# CITE-seq of murine bone marrow plasmacytoid dendritic cells and haematopoietic progenitor cells in sepsis

**DOI:** 10.64898/2026.07.15.738715

**Authors:** Johannes Korfhage, Elena Siakaeva, Mahamudul Hasan Bhuyan, Bettina Budeus, Dimitris Ttoouli, Tobias Lautwein, Stefanie B. Flohé, Stefanie Scheu

## Abstract

This dataset contains single-cell transcriptomic and surface protein profiles of bone marrow cells enriched for plasmacytoid dendritic cells (pDCs) and haematopoietic progenitor populations from septic and sham-operated mice. Sepsis was induced in BALB/c wild-type mice using cecal ligation and puncture (CLP) of moderate severity, with sham surgery as control. Bone marrow was collected 36 hours after surgery, depleted for selected lineage-positive cells by fluorescence-activated cell sorting, and processed using Cellular Indexing of Transcriptomes and Epitopes by Sequencing (CITE-seq). Sample demultiplexing was performed using hashtag oligonucleotides. The dataset includes gene expression matrices, antibody-derived tag counts, and barcode assignments for 13,725 cells, with a comparable distribution of sham and CLP samples across the total cell number. It may be used to study transcriptional states, surface marker expression, and phenotypic diversity of bone marrow cells in sepsis, to compare with other immunological and haematological single-cell datasets, or to train and benchmark computational tools for multimodal single-cell data analysis.

## Background & Summary

Sepsis is a life-threatening syndrome defined by organ dysfunction resulting from a dysregulated host response to infection^1^. In 2021, it globally affected 166 million people, with an estimated 21 million sepsis-related deaths, which represented almost 30% of total deaths worldwide and remains a major cause of morbidity and mortality^2^. The immune response in sepsis is characterized by an early “cytokine storm”, driven by the activation of the innate immune system, cytokine release, reactive oxygen species, endothelial activation, and the complement and coagulation systems, which may result in tissue damage, disseminated intravascular coagulation, and life-threatening organ failure. Patients who survive early hyperinflammation are at increased risk of secondary infections due to the concurrent development of immunosuppression. The development of immunosuppression during sepsis is associated with lymphocyte apoptosis, T cell exhaustion, induction of monocyte tolerance, reprogramming of dendritic cells (DC), impaired cytokine production, and expansion of regulatory T cells and myeloid-derived suppressor cells^3,4^. The pathomechanisms underlying the increased susceptibility to secondary bacterial infections in sepsis are not yet completely understood^5^. Multiple immune cell populations contribute to these processes, including pDCs, which are specialized in the rapid secretion of type I interferons in response to pathogen-derived nucleic acids^3,6-8^. DCs are key regulators of immunity but are highly vulnerable to sepsis-induced apoptosis and functional impairment, with loss evident in pDCs as well as conventional DCs (cDCs). These defects are associated with higher sepsis mortality and highlight the need for deeper molecular profiling of DC subsets^3^.

A widely used model to study human sepsis is “cecal ligation and puncture”, which induces septic peritonitis due to the release of intestinal bacteria^9,10^. Using this murine sepsis model, sepsis-activated pDCs have been shown to alter the function of DCs generated from the bone marrow, leading to increased IL-10 production; removal of these pDCs restored normal DC function^11^. These findings suggest that pDCs may influence DC differentiation in the bone marrow during sepsis through so far unknown mechanisms. To enable a detailed characterization of pDCs and DC progenitor cells in the bone marrow during sepsis, we generated a CITE-seq dataset from three septic and three control mice. Sepsis was induced by CLP while control mice underwent sham surgery. CLP involves ligation and perforation of the cecum that causes the release of gut content into the peritoneal cavity and subsequent systemic inflammation due to polymicrobial infection. The severity of sepsis in this model may be adjusted by the length of the ligation of the cecum and the size of the needle used to puncture the cecum. Here, we induced a moderate severity of sepsis that is associated with low mortality (20%) but increased susceptibility to secondary pulmonary infection^12^. Bone marrow cells were isolated 36 hours after surgery, the time point of maximal accumulation of activated pDCs in the bone marrow after CLP^11^.Lineage-positive cells except for pDCs were largely depleted from the bone marrow by FACS-sorting, resulting in the enrichment of pDCs and diverse haematopoietic progenitor cells. These enriched pDCs and progenitor cells were profiled for transcriptomic and surface protein expression using the 10x Genomics Chromium platform with Feature Barcoding technology. The dataset contains single-cell RNA sequencing (scRNA-seq) data, antibody-derived tag (ADT) measurements for cell surface markers, and hashtag oligonucleotide (HTO) information for sample demultiplexing.

This dataset will facilitate the exploration of transcriptional states, surface marker expression, and phenotypic heterogeneity of pDCs and DC precursors in murine sepsis. It can be integrated with other single-cell datasets to enable comparative analyses across disease models, time points, and experimental conditions. Potential applications include the investigation of sepsis-induced immune modulation, cross-species comparisons with human data, and the training or benchmarking of computational pipelines for multimodal single-cell data processing, clustering, and cell type annotation.

## Methods

### Animals

Female wild-type BALB/c mice (10-13 weeks old, 20.5-21.2 g) were obtained from Janvier Labs, Saint Berthevin Cedex, France. Mice were housed under specific pathogen-free (SPF) conditions with ad libitum access to standard rodent food and water at the animal facility of the University Hospital Essen. All animal procedures were approved by the North Rhine Westphalia State Agency for Nature, Environment and Consumer Protection, Germany (G1907/22 AZ81-02.04.2022.A214).

### Cecal ligation and puncture and sham surgery

Polymicrobial sepsis was induced using the CLP model, which recapitulates key features of human sepsis by releasing intestinal content into the peritoneal cavity and triggering a systemic inflammatory response as described before^11^. Sham surgery was performed as a negative control, following the same steps as the CLP procedure but without cecum manipulation (Fig. 1a)

**Figure 1.**
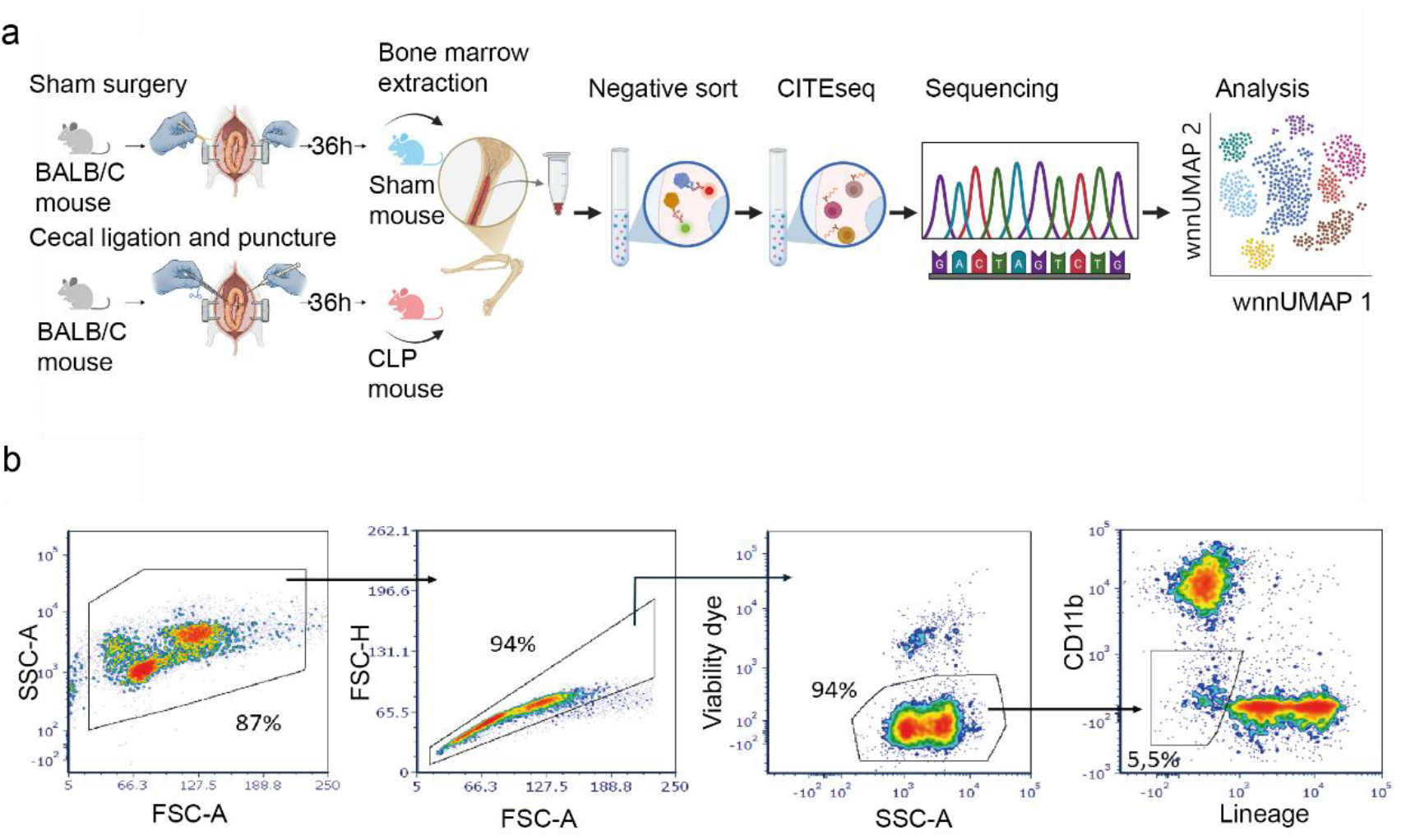
Experimental setup. (a) Three age and weight matched female wild-type BALB/c mice underwent sham and CLP surgery, respectively. Bone marrow was harvested from femora and tibiae 36 h post-surgery. After FACS sorting, the enriched cell populations were subjected to CITE-seq antibody staining. Single-cell libraries were generated using 10x Genomics Chromium Controller and sequenced on an Illumina NextSeq 2000 platform prior to downstream bioinformatics analysis. (b) The gating strategy for excluding non-target cell populations by fluorescence-activated cell sorting (FACS).

For CLP, mice were grouped by body weight and anesthetized via intraperitoneal injection of ketamine (100 mg/kg body weight; WDT, Garbsen) and xylazine (10 mg/kg body weight; Rompun, Bayer) adjusted in 0.9% NaCl (Fresenius Kabi) to a final injection volume of 200 μl. Adequate anaesthesia was confirmed by absence of the interdigital pinch reflex. In cases of insufficient anaesthesia, additional doses of ketamine and xylazine were administered. Bepanthen ophthalmic ointment (Bayer) was applied to the eyes to prevent them from drying, and mice were placed on a 37 °C heating plate. After abdominal skin disinfection, a ∼1.5 cm midline laparotomy was performed under aseptic conditions. The cecum was exposed and ligated at 35% of its length from the distal tip with a polyglycolic acid (PGA) surgical thread (5-0, Resorba). Subsequently, the ligated part of the cecum was punctured once with a 27-gauge needle (Sterican, B. Braun), and ∼2 μl of fecal material was gently extruded. The cecum was returned to the peritoneal cavity, and 1 ml of warm (37°C) sterile 0.9% NaCl was administered into the peritoneal cavity for resuscitation. The incision was closed in two layers: peritoneum – with PGA 5-0 surgical thread; skin – with ETHICON PROLENE surgical thread (5-0; Johnson & Johnson). Finally, Buprenorphine (0.1 mg/kg in 30 μl 0.9% NaCl; Buprenovet, Bayer) was administered subcutaneously for pain relief. The animals did not receive any antibiotics. After surgery, mice were regularly monitored, and their condition was assessed to allow early identification of humane endpoints.

### Bone marrow isolation

Animals were sacrificed by cervical dislocation. Skin was removed via an incision from the lower pelvis to the abdomen, and muscles and connective tissue were dissected to detach the hind limbs. Feet and residual tissue were removed, bones (femur and tibia) were separated from each other, disinfected with 70% ethanol, and stored in medium (Very low endotoxin (VLE) RPMI 1640 (BioSell, BS.FG1415) supplemented with 8% heat-inactivated FCS (Gibco), 10 mM Hepes Buffer (ThermoFisher Scientific), 100 IU/ml penicillin (Sigma-Aldrich), 0.02 mg/ml gentamicin (Sigma-Aldrich) and 0.05 mM of 2-mercaptoethanol (Sigma-Aldrich)) on ice until proceeding with the cell isolation. Bones were briefly transferred through two portions of 70% ethanol and then washed in fresh sterile PBS. Both ends of the tibiae were cut, while femora were cut in the middle into two parts. The bone marrow was flushed out with cold medium using a syringe with a 27-gauge needle and collected into a sterile plate for subsequent processing, then resuspended with a 100 µl pipette tip and transferred to a Falcon tube via a 30 µm cell strainer. Cells were pelleted (6 min, 300 × g, 6 °C). Red blood cells were lysed using 2 ml of Erylyse buffer for 2 min before adding 5 ml medium. After lysis, cells were pelleted again, resuspended in fresh medium and counted using a Neubauer counting chamber.

### CITE-seq sample preparation

Selected lin^-^ and live bone marrow cells were enriched by negative FACS sorting against the surface markers CD11b, CD19, CD49, TCRβ, and Ter119 and Ef780 for dead cell exclusion after Fc receptors were blocked to minimize nonspecific antibody binding and improve staining specificity. (Fig. 1b). For sample demultiplexing, cells from the six individual animals were incubated according to the manufactures instructions with hashtag oligonucleotide (HTO) antibodies (BioLegend; 1:100) for 30 min at 4 °C, assigned as follows: CLP1-HTO1, CLP2-HTO2, CLP3-HTO3, Sham4-HTO4, Sham5-HTO5, Sham6-HTO6. Unbound HTOs were removed by three washing steps with FACS buffer (PBS containing 2% FCS, 2 mM EDTA). The respective Sham and CLP samples were pooled, and Fc-block treatment was repeated (10 min at 4 °C). The TotalSeq-A cocktail (BioLegend) was rehydrated according to the manufacturer’s instructions and supplemented with spike-in antibodies against CD80 (1:20), Ly6D (1:20), CD135 (1:5), CD34 (1:10), and cKit (1:40) (all BioLegend). A complete list of antibodies and their total ADT UMI counts per marker is provided in Supplementary Figure 1. Spike-in antibody–oligonucleotide conjugates were titrated prior to CITE-seq to determine the optimal staining concentration as follows: Bone marrow cells were stained in parallel with serial dilutions of the antibody (1:10, 1:20, 1:50, 1:100, 1:200), maintaining constant cell numbers and staining volume. The optimal antibody concentration was determined by flow cytometry as the highest dilution yielding maximal separation between positive and negative populations (greatest signal-to-noise ratio) without increased background in negative cells; for subsequent CITE-seq experiments, the next higher dilution was selected to minimize nonspecific binding. One microliter of each dilution was added to the cocktail. After centrifugation to remove aggregates (10,000 × g, 30 s; followed by 14,000 × g, 10 min, 4 °C), 25 µl of antibody mix was added to 25 µl Fc-blocked cells and incubated for 30 min at 4 °C. Cells were washed, re-sorted for viability, and 150,000 live cells were collected. Cells were washed in resuspension buffer (0.04% BSA in PBS), adjusted to 1,000 cells/µl, and viability was confirmed microscopically (Fig. 1).

### Single-cell library preparation and sequencing

Approximately 16,000 cells were processed using the Chromium Controller (10x Genomics, Pleasanton) and the Chromium Single Cell 3′ NextGEM Reagent Kit v3.1 with Feature Barcoding technology, according to the manufacturer’s protocol. Sequencing was performed on a NextSeq 2000 platform (Illumina) with a mean depth of 57,000 reads per cell for gene expression libraries, ∼10,000 reads per cell for ADT libraries, and ∼5,000 reads per cell for HTO libraries.

### Processing of 10x Genomics single-cell data

Raw sequencing data were processed using Cell Ranger software (v7.2, 10x Genomics). BCL files were demultiplexed into FASTQ files using the cellranger mkfastq pipeline. Alignment to the mm10 reference genome (GRCm38) and UMI counting were performed using the cellranger count pipeline to generate gene–barcode matrices.

### CITE-seq data processing and quality control

Data analysis was performed in R (v4.3.3; R Core Team) using RStudio (Posit) and Seurat (v5.1.0; RRID:SCR_016341). Required packages included ggplot2 (RRID:SCR_014601), patchwork and dplyr (RRID:SCR_019163). Filtered barcode and feature matrices were imported, and HTOs were renamed according to their sample IDs. HTO demultiplexing was used to classify cells as negative, singlet, or doublet; only singlets were retained for downstream analysis. Seurat objects were created for gene expression, HTO, and ADT data. Low-quality cells were filtered out. Cells with a mt (mitochondrial transcripts) percentage >10%, nFeature_RNA >300 and nCount_RNA <75,000 were excluded. Centered log-ratio (CLR) normalization was applied to HTO and ADT assays. ADT and HTO data were processed, and the wnn UMAP was plotted (Fig. 1). For ADT quality control, isotype control features were identified and pooled to define the background signal distribution. For each non-isotype antibody, the area under the receiver operating characteristic (ROC) curve was calculated by comparing its normalized signal distribution with the pooled isotype control distribution. In addition, a stain index was calculated for each antibody as the difference between the median signal within its highest 10 percent of measurements and the mean isotype signal, divided by twice the standard deviation of the isotype signal. Antibody features with an area under the curve <0.50 and a stain index <0.30 were considered insufficiently separated from background and were excluded. All isotype control features were removed before downstream analysis. The retained ADT features were subsequently normalized and scaled before principal component analysis and weighted nearest neighbour integration.

### Data Records

The raw CITE-seq data generated for this study have been deposited in the European Nucleotide Archive (ENA) under the study accession PRJEB101194 (secondary accession ERP182617), entitled “CITE-seq of murine bone marrow dendritic cells and precursors in sepsis.” (https://www.ebi.ac.uk/ena/browser/view/PRJEB101194). The dataset comprises three library components, gene expression (GE), antibody-derived tags (ADT), and hashtag oligos (HTO), each represented as an individual Sample, Experiment, and Run within the ENA submission framework.

Three sample records were created, corresponding to the three CITE-seq modalities:

- ERS28029860 (SAMEA120615168), representing the ADT library
- ERS28029861 (SAMEA120615169), representing the GE library
- ERS28029862 (SAMEA120615170), representing the HTO library

Each sample record is linked to a dedicated Experiment record describing the library preparation and sequencing configuration (ERX15359314, ERX15359315, and ERX15359316, respectively). The resulting raw data files are stored as three Run accessions, ERR15964205 (ADT), ERR15964206 (GE), and ERR15964207 (HTO), generated on an Illumina NextSeq 2000 platform. Each Run contains one paired-end FASTQ file set.

The ENA organizes the deposited data using its standard hierarchical structure (Study → Sample → Experiment → Run). Within this structure, the raw FASTQ files are located in Run-specific directories, and all associated metadata, including the submission spreadsheet and XML descriptors for Samples, Experiments, and Runs, is stored in a dedicated metadata folder.

### Technical Validation

Sequencing of bone marrow–derived pDCs and diverse haematopoietic progenitor cells yielded a total of 13,725 high-quality single cells. Each cell was sequenced to a mean depth of 57,189 reads, with a median of 14,200 unique molecular identifiers (UMIs) and 4,297 detected genes per cell (Fig. 2a). Across all libraries, 26,138 genes were detected in total. Read quality was high, with Q30 base scores exceeding 91% for RNA reads, 94% for barcodes, and 94% for UMIs. ADTs were also of high quality, with valid barcodes in 98.7% of reads, a sequencing saturation of 81.4%, and a median of 1,322 antibody UMIs per cell (Fig. 2b). The majority of antibody reads (63.3%) were assigned to recognized antibody barcodes, with 90.4% of sequencing reads corresponding to antibody features. Mapping efficiency was robust: 95.8% of reads aligned to the genome, with 92.6% mapping confidently. Among these, 59.0% mapped to exonic regions, 30.3% to intronic regions, and 3.4% to intergenic regions. Approximately 76% of reads mapped confidently to the transcriptome, and only 13.1% were antisense to annotated genes. These statistics support the technical integrity of the dataset. ADT quality control showed that most antibody features were clearly separated from the isotype control background. Five antibody features, CD23, Ly49D, CD19, TCR B chain, and TER 119, were insufficiently separated from background and were therefore excluded. All remaining antibody features showed clear separation from the background and were retained for downstream analysis. Cells were filtered to exclude potential low-quality cells based on gene count, UMI count, and proportion of mitochondrial transcripts (see Methods for thresholds). Additionally, HTO demultiplexing was part of the core QC to remove multiplets and ambiguous cells based on HTOs before downstream analysis (Fig. 3). This ensured that downstream analyses were restricted to cells with reliable transcriptomic and proteomic profiles. Technical consistency was observed across experimental conditions. In addition, concordance between transcriptomic and proteomic modalities was observed for canonical lineage-defining markers such as CD317 (pDCs), CD25 (ILC2), and CD200R3 (BaPs), where RNA expression and ADT signals exhibited highly similar spatial distributions ^13-16^. Weighted nearest neighbour (WNN)-based UMAP projections showed that the RNA expression of key surface markers, including Bst2 (CD317), Il2ra (CD25), and Cd200r3, was highly enriched in defined cellular clusters. Importantly, the corresponding ADT signals for CD317, CD25, and CD200R3 exhibited nearly identical spatial distributions (Fig. 4). This close agreement between RNA and protein measurements of markers with established correlations for these modalities confirms the accuracy of the CITE-seq assay, validates the integrity of multimodal integration, and supports the reliability of downstream analyses based on combined transcriptomic and proteomic information. Although this agreement was pronounced for established markers, the multimodal framework also enabled the detection of instances in which transcript and protein levels were not fully aligned, potentially reflecting post-transcriptional regulation or context-dependent expression dynamics. Quality metrics, including distribution between conditions such as sham and CLP (Fig. 5a) for single mice, and conditions and cells from those mice were comparable between sham and septic mice, supporting the reproducibility of the dataset (see Fig. 5b for CLP conditions and Fig. 5c for sham condition). Together, these metrics demonstrate that the dataset is of high technical quality, with sufficient sequencing depth, robust read mapping, and biologically coherent multimodal profiles suitable for downstream analysis and reuse.

**Figure 2.**
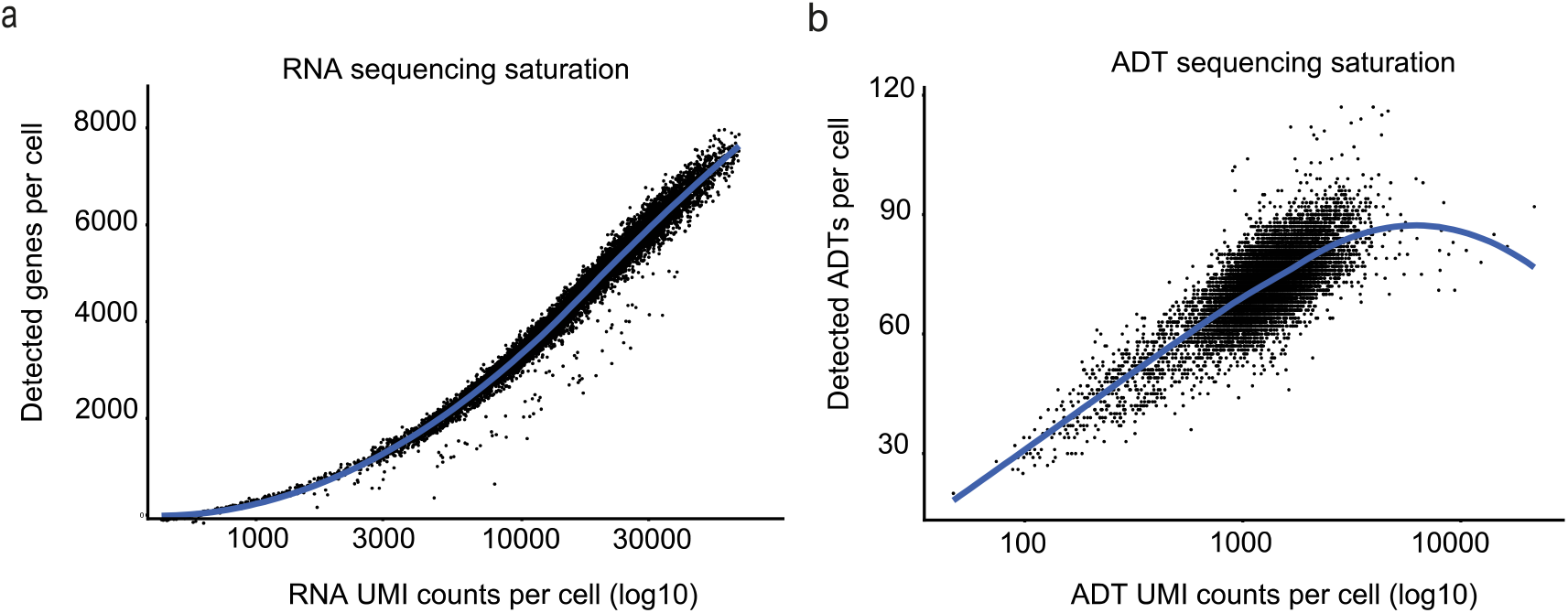
RNA and ADT sequencing saturation curve: (a) For each cell, the number of detected genes was plotted against the total RNA UMI counts (log10 scale). The resulting curve shows a steep incline in the number of detected genes at low UMI count, followed by a progressive decline in slope at higher UMI count, indicating diminishing returns in gene detection with increasing sequencing depth. The median sequencing depth of 14,200 RNA UMIs per cell lies within this region of reduced gain, suggesting that most transcriptomes were captured close to saturation. (b) Number of detected antibody-derived-tags (ADTs) as a function of ADT UMI counts per cell. With increasing sequencing depth, the number of detected ADTs rises initially and subsequently approaches a plateau, consistent with a Heidelberger-type saturation curve. This behaviour indicates that most antibody targets were captured near saturation and that additional sequencing would yield diminishing returns in terms of newly detected protein features. Together, both plots demonstrate adequate sequencing depth for reliable gene expression quantification across the dataset and confirm high-quality ADT detection that supports the sustainability of the dataset for robust multimodal integration.

**Figure 3.**
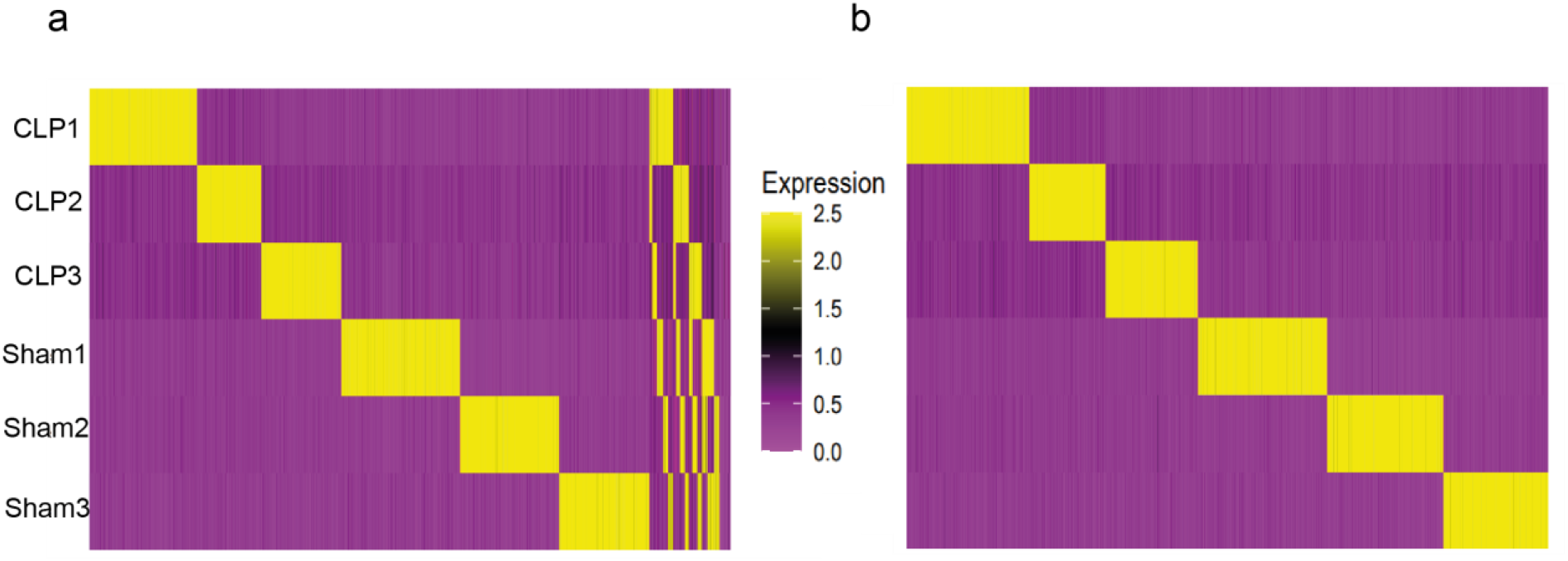
HTO Demultiplexing Quality Control: (a) The heatmap visualizes normalized HTO signal intensities across all barcoded droplets, showing six sharply separated hashtag patterns corresponding to the experimental samples CLP1–3 and sham1–3. Strong diagonal blocks indicate robust sample-specific HTO enrichment, while off-target signal remains low. This confirms successful sample multiplexing before removal of doublets and negatives. (b) The same heatmap as in (a) is shown after HTO Demux classification and filtering. The heatmap shows well-defined and mutually exclusive HTO signals for all six samples. The absence of mixed or ambiguous blocks indicates effective identification and removal of doublets and unclassified droplets, ensuring high sample purity for downstream analysis.

**Figure 4.**
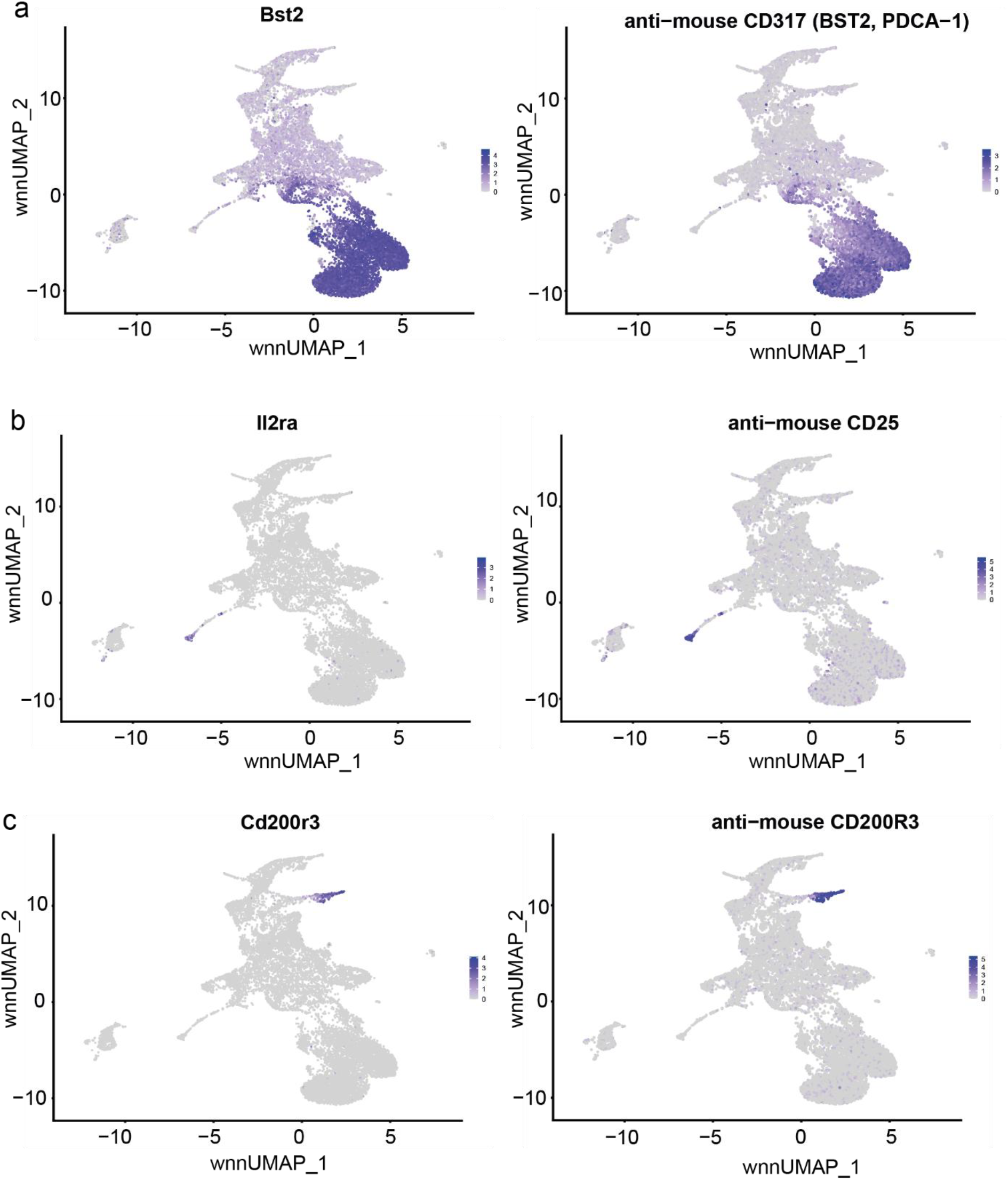
Concordance between RNA and protein modalities: A wnnUMAP is coloured by RNA expression of Bst2 (CD317) (a, left column), Il2ra (CD25) (b, left column) and Cd200r3 (c, left column). The respective genes are strongly enriched in defined clusters. Right column shows ADT signals for anti-mouse CD317 (a, right column), anti-mouse CD25 (b, right column), and anti-mouse CD200R3 (c, right column). Nearly identical spatial patterns of the RNA- and ADT-signals validate the accuracy of the multimodal CITE-seq measurements and support reliable downstream analyses.

**Figure 5.**
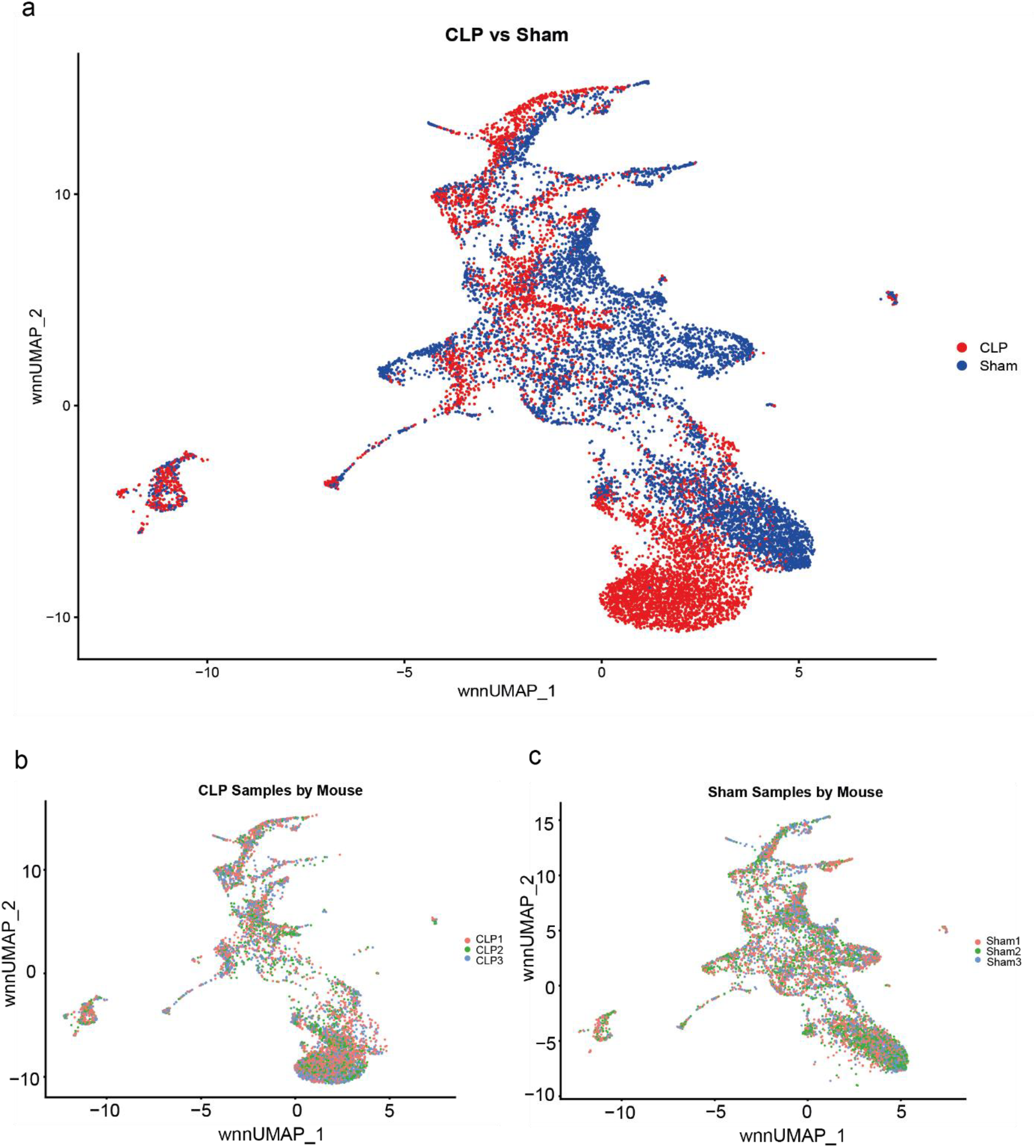
Clustering and batch effect assessment: (a) Condition-level mapping shows a partial overlap between CLP and sham cells, indicating shared myeloid populations, while also highlighting condition-specific shifts in cluster abundance. Importantly, there is no global condition-driven embedding artifact, further supporting low batch contributions. (b) CLP samples from three independent mice map uniformly across the UMAP, with no replicate-specific clustering. This demonstrates strong biological reproducibility and minimal batch effects among specific animals. (c) Sham replicates are equally in the UMAP space, confirming that control animals also lack batch-driven segregation. Together, these plots show that sample identity does not drive clustering structure, supporting the robustness of the dataset.

## Data availability

All raw CITE-seq data generated in this study have been deposited in the European Nucleotide Archive (ENA) under the study accession **PRJEB101194** (https://www.ebi.ac.uk/ena/browser/view/PRJEB101194). The submission includes separate Sample, Experiment, and Run records for the gene expression (GE), antibody-derived tag (ADT), and hashtag oligo (HTO) libraries. All associated metadata describing sample characteristics, library preparation, and sequencing parameters are provided within the ENA record.

## Code Availability

All custom R scripts used for pre-processing and analysis are available at GitHub (https://github.com/Johannes-Korfhage/CITEseq-sepsis-code-1).

## Author Contribution

JK and ES contributed equally to this work. JK and ES performed the experiments, including animal work, and conducted the bioinformatics analyses. JK prepared the figures and wrote the original draft of the manuscript. EL contributed to data interpretation and manuscript revision. TL performed the sequencing and contributed to data generation and processing. MB assisted with experimental design and data analysis. BB and DT provided support for bioinformatics analyses and code development. SF and SS supervised the study and served as principal investigators of the participating laboratories. All authors reviewed and approved the final manuscript.

## Acknowledgements

We thank all members of the participating laboratories for helpful discussions and technical support. We acknowledge the support of the animal facility staff for excellent animal care and assistance with mouse experiments. We thank the sequencing and genomics core facility for their support with library preparation and sequencing. We are grateful to the bioinformatics and high-performance computing infrastructure for providing computational resources.

Stefanie Scheu was supported by the Deutsche Forschungsgemeinschaft (DFG, German Research Foundation) projects SCHE692/6–1, SCHE692/8–1(465743247), and DFG-

270650915/GRK2158 and the Manchot Graduate School ‘Molecules of Infection IV’.

Stefanie Flohé was supported by the Deutsche Forschungsgemeinschaft (DFG, German Research Foundation) projects FL 391/6-1 (465743247).

## Generative AI Statement

The authors used DeepL (DeepL SE, Cologne, Germany) and ChatGPT (OpenAI) exclusively for language editing, improvement of manuscript readability, and assistance with code optimization and documentation. All scientific content, data interpretation, analyses, and conclusions were developed and verified by the authors, who take full responsibility for the final manuscript.

**Supplementary Figure 1.**
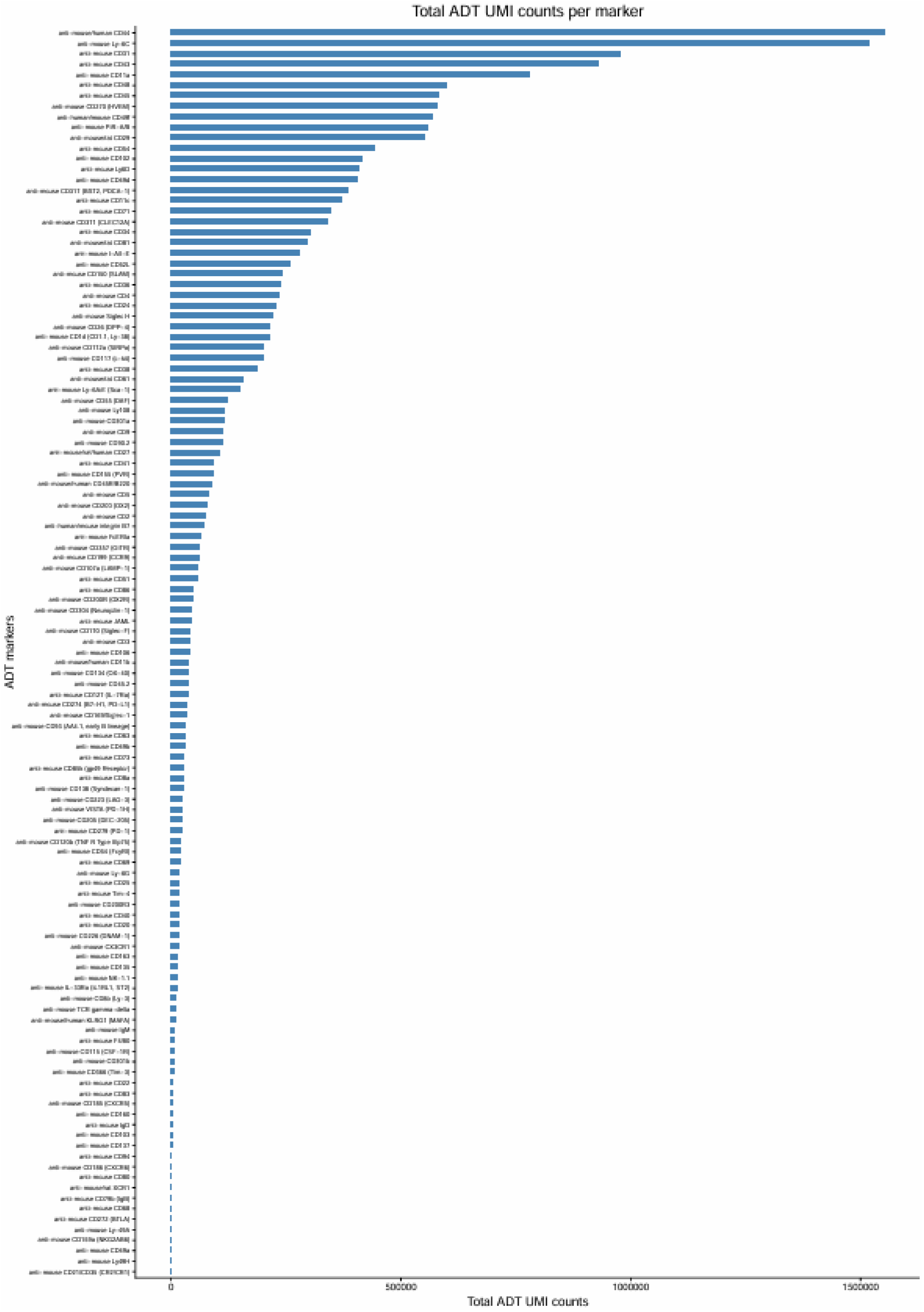
Distribution of total ADT UMI counts across protein markers. Total antibody-derived tag (ADT) unique molecular identifier (UMI) counts for each protein marker measured by CITE-seq, summed across all cells after quality control. Markers are ordered by decreasing total ADT UMI counts. This plot summarizes the overall detection levels of the ADT panel and serves as a quality control metric for protein expression profiling.

